# Responses of soil physical, chemical and biochemical properties to short-term impact of tillage in an alfisol under maize (*Zea mays* L.) cultivation

**DOI:** 10.1101/542944

**Authors:** Ajoke Adegaye, Solomon Adejoro, Segun Oladele, Daniel Arotupin, Stephen Ojeniyi

## Abstract

A short-term field and laboratory experiment was conducted to assess the impact of four traditional tillage methods on the physical, chemical and biochemical properties of a sandy clay loam alfisol under maize (*Zea mays* L.) cultivation following a fallow period of five years. Treatments included slash only (SO), slash +burn (SB), slash+ ridge (SR) and herbicide (glyphosate) application (HA) at the recommended rate of 2 L/ha^-1^. Results from the statistical analysis of data from this study showed inconsistent trends of tillage treatments on soil physical properties. However, slash +burn (SB) tillage significantly (p < 0.05) enhanced chemical properties such as the soil pH, available P, exchangeable K ^+^, Ca^2+^, and Mg^2+^ compared to their respective values before treatment application in the two years under study. Amongst the treatments, slash + burn (SB) tillage further exerted the most significant (P ≤ 0.05) effect on urease, L-asparaginase, L-glutaminase, dehydrogenase, acid and alkaline phosphatase activities, but recorded the least values for amidase and β-glucosidase activities in soil. Results from this study therefore, suggest that SB tillage treatment was beneficial to the soil environment as it has proven to be more promising and effective for enhancing the selected soil quality indicators on the soil type due to quick mineralization and release of bound nutrients present in the soil and litter.

## 1. Introduction

The activities of soil enzymes have been predominantly used in most fields as biological indicators of soil quality to rapidly discover management-induced alterations in microbial activity and/or SOM composition (Kandeler et al., 1999), stabilizing the soil structure and other life processes of soil microbes, play key ecological functions in organic matter decomposition, mineralization and nutrient cycling processes (Dick 1994; Kandeler et al., 2006). Soil enzyme activity measurement, due to their ease of quantification, has long been connected with soil fertility evaluation and sustenance in agrarian ecosystems with a high turnover of organic compounds.

In general, agricultural induced soil disturbance, as one of the many factors affecting soil microbial and biochemical properties, has been a major cause of OM losses in fields (Bayer et al., 2001). The SOM serves as a major source and sink of nutrients in soil, regulates all soil microbial processes and activity and enhances the soil’s physical and chemical properties. However, its loss usually leads to decline in soil fertility status; hence, crop productivity levels (Gregorich et al., 1994).

Several researches have been carried out to investigate the effects of conservative tillage methods on the physical, chemical and microbiological soil properties (Staddon et al., 1998; Bausenwein et al., 2008; Jin et al., 2009). These tillage practices are principally envisioned to limit erosion problems, conserve or improve SOM composition, enhance other SOM-influenced soil properties and consecutively improve crop production. Further researches have also suggested the advantages of conservative tillage practices on enzyme activities in subtropical soils (Roldán et al., 2005a, 2005b) as well as under the semi-arid Mediterranean and temperate climate (Kandeler et al., 1999; Riffaldi et al., 2002). However, available information on the effects of these traditional tillage methods on the soil physical, chemical and biochemical quality indices on the Nigerian tropical soils is scanty.

Therefore, this work aimed to assess the impacts of tillage treatments on (1) soil physical properties including soil bulk density, total porosity, water holding capacity and soil temperature; (2) soil chemical properties and (3) activities of urease, amidase, L-asparaginase, L-glutaminase, β-glucosidase, dehydrogenase, acid and alkaline phosphatases in soils.

## 2. Materials and methods

### 2.1 Experimental site characteristics

The present study was conducted in two years at the Teaching and Research Farm of the Federal University of Technology, Akure (7°20 N, 5°30E) Ondo State, Nigeria. The site was characterized by a semi-arid to humid climate with a mean annual temperature of 26°C, relative humidity of about 15% and average annual precipitation of about 1400 mm, most of which occurs from May to September. The topography of the area is flat and slope does not exceed 1%. The understory vegetation of the experimental field after about five years of fallow consisted predominantly of some weed species such as *Impereta cylinderica, Chromolaena odorata, Talinum triangulare, Ageratum conizoides, Penisetum purpureum, Gliricidia sepium, Euphorbia heterophylla, Sida acuta, Cynodon dactylon* and *Panicum maximum.* The soil at the site is classified as Typic Paleustalf Alfisol (Soil Survey Staff, 2014), with clay, sand and silt contents of 22%, 67% and 11% respectively (Tel, 1984). The average pH (1:2.5 soil–H_2_O; McLean, 1982); available P using the Bray-I method (Bray and Kurtz, 1945); SOC using the Walkley and Black (1934) method; total N content by the Kjeldahl digestion (Nelson and Sommers, 1982); exchangeable cations and CEC using ammonium acetate method (Black, 1965); bulk density, total porosity, water holding capacity and temperature were all determined as well.

### 2.2 Experimental set-up and procedure

The total field size was 15 m by 12.5 m (187.5 m^2^) with each experimental unit measuring 3.0 × 3.5 m with 1 m alley. The experiment was laid out in a Randomized Complete Block Design (RCBD), involving four tillage treatments with three replications per block. Treatments included slash only (SO), slash and burn (SB), slash and ridge (SR), and herbicide (glyphosate) application (HA) at the field recommended rate of 2.0 l/ha. The four tillage treatments were established and two maize seeds were sown initially at a spacing of 75cm × 35cm inter and intra row spacing respectively, but was later thinned to one plant stand 2 weeks later. The improved maize variety was obtained from the International Institute of Tropical Agriculture (IITA), Ibadan in Nigeria. Following crop emergence, subsequent weed outgrowth on the field was cleared manually to prevent any form of chemical interaction with biochemical analysis.

### 2.3 Soil sampling and pre-treatment

Five soil sample cores (1.5 cm diameter) were randomly collected on each experimental unit at a depth of 0-15 cm during the maize growing season between April and September, 2017 and 2018 respectively and analysed for its physical, chemical and biochemical properties. Soil samples collected for physical and chemical analyses were bulked, air-dried and sieved (2 mm), while samples for biochemical analyses were placed in plastic bags and kept on ice while being transported back to the laboratory, mixed to form one composite sample/treatment, sieved (2 mm), stored at 4°C, and analyzed within 24 h of collection.

### 2.4 Assay for enzyme activities

The activities of urease, amidase, L-asparaginase, L-glutaminase, β-glucosidase, acid and alkaline phosphatases were all assayed using the described procedures of Tabatabai (1994). Nitrogen cycling enzyme (urease, amidase, L-asparaginase, L-glutaminase) activities were determined by measuring the NH^4+;^–N released in the hydrolysis reaction and results expressed as mg NH^4+;^–N g^-1^ dwt 2h^-1^. The activities of β-glucosidase, acid and alkaline phosphatases were determined by measuring *p*-nitrophenol concentration by spectrophotometry at 400 nm and results expressed as µg ρ-nitrolphenol g^-1^ dwt h^-1^. Dehydrogenase activity (DHA) was determined according to the standard method of Casida et al. (1964). The Triphenylformazan (TPF) formed after incubation was extracted with 10 mL of methanol and measured spectrophotometrically at 485 nm. Dehydrogenase results were expressed as µg TPF g^-1^ dwt h^-1^. Each sample was performed in triplicates and controls followed the same procedure, but with the exception that the substrate was added after the incubation and addition of reaction stopper solutions. The assay conditions are further summarized in Table 1 and results communicated on a moisture-free basis. Moisture contents of samples from each tillage treatment were determined by drying the soil samples at 105°C for 24 h. All laboratory experiments were conducted at the Soil Science laboratory of the Department of Crop, Soil and Pest Management and the Biochemical laboratory of the Department of Biochemistry, the Federal University of Technology, Akure (FUTA), Ondo State, Nigeria.

**Table 1:** Assay conditions used to determine the activities of the soil enzymes

| Enzyme activity | EC | Substrate used | Buffer | pH | Temperature<br>(°C) | Time (h) |
| --- | --- | --- | --- | --- | --- | --- |
| Urease | EC 3.5.1.5 | Urea | THAM | 9.0 | 37 | 2 |
| amidase | EC 3.5.1.4 | formamide | THAM | 8.5 | 37 | 2 |
| L-asparaginase | EC 3.5.1.1 | L-asparagine | THAM | 10 | 37 | 2 |
| L-glutaminase | EC 3.5.1.2 | L-glutamine | THAM | 10 | 37 | 2 |
| β- glucosidase | EC 3.2.1.21 | ρ-nitrophenyl-β-D-glucoside (PNG,<br>0.05M) | MUB,<br>THAM | 6.0, 12 | 37 | 1 |
| dehydrogenase | EC 1.1 | 2,3,5-Triphenyltetrazolium<br>chloride (TTC, 3%) |  |  | 37 | 24 |
| acid phosphatase | EC 3.1.3.1 | ρ-nitrophenyl<br>phosphate disodium<br>(ρNPP, 0.05 M) | MUB | 6.5 | 37 | 1 |
| alkaline<br>Phosphatase | EC 3.1.3.2 | ρ-nitrophenyl<br>phosphate disodium<br>(ρNPP, 0.115M) | MUB | 11 | 37 | 1 |

\*THAM = tris-hydroxymethyl aminomethane, MUB = modified universal buffer

### 2.5 Statistical analysis

Statistical analysis of data collected was conducted using the one way analysis of variance (ANOVA) procedure in MINITAB Statistical Software 17th edition (MINITAB Inc., 2007). The significant difference was compared using Tukey test at p < 0.05 to indicate significant differences among the treatment means. Standard errors and means were estimated to construct graphs using Microsoft excel 2007.

## 3 Results

### 3.1 Soil physical properties

Table 2 shows that significant differences (P ≤ 0.05) exist in the effect of tillage treatments on soil bulk density (BD), total porosity (TP), water holding capacity (WHC) and temperature (T°C) in both years. In 2017, slash only gave the highest BD increase of 11.5%, least significant decreases in TP and WHC of −49% and −39% respectively, compared to their respective mean values before experiment. Slash and ridge (SR) had the least significant decrease of −6% for BD which resulted in the highest and only significant increase of 3% and 11% for TP and WHC of the soil respectively, while other tillage treatments decreased in comparison with their respective mean values before experiment. Highest and significant increase of about 44% in soil T°C was observed with SB treatment. In 2018, SO tillage treatment gave the highest value of 5% for BD, however, SR recorded a decreased value of −26% compared to their respective mean values before experiment. Also, SR had increased values of 12% and 8% for TP and WHC respectively although, not significantly different from those of SB (10%, 8%) and HA (5%, 13%) tillage treatments. Significant increase in soil T°C of about 34% was again observed with SB tillage plots but, no noticeable significant differences were recorded with other tillage treatments after treatment application.

**Table 2:** Effects of tillage methods on soil physical properties (0-15 cm) before and after tillage treatment application in 2017 and 2018 cropping trials.

| Soil physical properties | Bulk density (g/cm<sup>3</sup>) | Total porosity (%) | Water holding capacity (%) | Temperature (°C) at 10 cm depth |
| --- | --- | --- | --- | --- |
| **2017** |  |  |  |  |
| Before Exp. | 1.39b | 38.2a | 27.5ab | 27.33b |
| Slash + Burn | 1.42ab | 32.1ab | 22.2ab | 48.5a |
| Slash only | 1.57a | 25.6b | 19.8b | 32.7ab |
| Slash + Ridge | 1.31b | 39.2a | 30.9a | 30.8ab |
| Herbicide | 1.45ab | 30.7ab | 24.6ab | 32.9ab |
| **2018** |  |  |  |  |
| Before Exp. | 1.44ab | 31.5a | 32.4ab | 27.71b |
| Slash + Burn | 1.43ab | 34.8a | 35.2a | 41.7a |
| Slash only | 1.52a | 27.2b | 30.3b | 30.7ab |
| Slash + Ridge | 1.35b | 35.7a | 35.4a | 31.8ab |
| Herbicide | 1.45ab | 32.2a | 38.8a | 29.6ab |

Mean values in the same column followed by different letters in each year are significantly different from one another based on Tukey's test at $p \leq 0.05$ .

### 3.2 Soil chemical properties

Significant differences (P ≤ 0.05) were recorded on the soil chemical properties before and after experiments in both years (Table 3). When compared with their respective mean values before experiment in 2017, slash + burn (SB) increased the soil pH by 11%, K^+^ (25%), Ca^2+^ (7%) and Mg^2+^ (6%). Slash + ridge (SR) increased K^+^ by 26%. Only herbicide application (HA) gave an increase of 4% each for both OC and OM. in this study, there were significant reductions in the values of OC, OM, total N and available P after treatment application, while Na^2+^ content remained unchanged across the tillage treatments. Similar trend were obtained in 2018, slash + burn increased the soil pH by 18%, available P (59%), exchangeable K^+^ (64%), Ca^2+^ (87%) and Mg^2+^ (86%). Herbicide application (HA) increased the soil pH by 8%, exchangeable K^+^ (3%), Ca^2+^ (59%) and Mg^2+^ (77%). Therefore, there were increases in soil pH, available P, exchangeable K^+^, Ca^2+^ and Mg^2+^, while decrease in OC, OM, TN and Na^+^ were observed after treatment application across the tillage treatments.

**Table 3:** Pre- and post-soil chemical properties (0-15 cm) as affected by tillage treatments in 2017 and 2018 cropping trials.

| Soil chemical properties | pH (1:2 H<sub>2</sub>O) | OC | OM | Total N | Avail. P (mg kg<sup>-1</sup>) | Exchangeable cations |  |  |  |
| --- | --- | --- | --- | --- | --- | --- | --- | --- | --- |
|  |  |  |  |  |  | K<sup>+</sup> | Na<sup>+</sup> | Ca<sup>2+</sup> | Mg<sup>2+</sup> |
|  |  |  |  |  |  | (cmol kg<sup>-1</sup>) |  |  |  |
| 2017 |  |  |  |  |  |  |  |  |  |
| Before Exp. | 5.45b | 2.50ab | 4.31a | 1.25a | 3.08a | 1.23b | 1.04a | 2.97ab | 1.03a |
| Slash + Burn | 6.10a | 2.42ab | 4.17ab | 0.26b | 3.01a | 1.64a | 1.04a | 3.20a | 1.10a |
| Slash only | 5.60ab | 2.12b | 3.65b | 0.31b | 2.77a | 1.31ab | 1.09a | 2.13b | 0.77b |
| Slash + Ridge | 5.20c | 2.20b | 3.79b | 0.35b | 2.85a | 1.67a | 1.04a | 2.13b | 0.87ab |
| Herbicide | 5.68ab | 2.60a | 4.48a | 0.30b | 2.98a | 1.23b | 1.04a | 2.97ab | 0.80ab |
| 2018 |  |  |  |  |  |  |  |  |  |
| Before Exp. | 5.21c | 2.37a | 4.09a | 1.27a | 1.23bc | 0.36b | 1.22a | 1.16c | 0.41c |
| Slash + Burn | 6.34a | 1.55b | 1.12b | 0.10b | 2.98a | 0.99a | 0.43b | 8.77a | 2.97a |
| Slash only | 5.42b | 1.90ab | 1.38b | 0.22b | 1.57b | 0.95a | 0.27c | 4.30b | 1.57b |
| Slash + Ridge | 5.41b | 1.56b | 1.13b | 0.18b | 1.48b | 0.22b | 0.51b | 4.07b | 2.77a |
| Herbicide | 5.64b | 1.53b | 1.11b | 0.20b | 0.20c | 0.37b | 0.22c | 2.83bc | 1.77b |

### 3.3 N-cycling enzyme activities

Table 4 shows the effects of tillage methods on N-cycling enzyme activities in the soil. This study showed that tillage methods significantly (P ≤ 0.05) affected the soil L-glutaminase, L-asparaginase, urease and amidase activities. Slash + burn had the highest soil L-glutaminase (54.54 mg NH_4_-N/g dwt/2h), urease (45.21 mg NH_4_-N/g dwt/2h) and L-asparaginase (33.25 mg NH_4_-N/g dwt/2h) activities compared to other tillage methods. The lowest L-glutaminase activity was recorded with slash only (2.33 mg NH_4_-N/g dwt/2h), while lowest urease (4.96 mg NH_4_-N/g dwt/2h) and L-asparaginase (3.50 mg NH_4_-N/g dwt/2h) activities were recorded with slash + ridge. Herbicide application (63.29 mg NH_4_-N/g dwt/2h) had the highest amidase activity which does not differ significantly from slash only (49.29 mg NH_4_-N/g dwt/2h) but, differed significantly from slash + burn (−29.75 mg NH_4_-N/g dwt/2h) and slash + ridge (−18.96 mg NH_4_-N/g dwt/2h) plots.

**Table 4:** Effects of tillage methods on N-cycling enzyme activities

| Treatment | L-Glutaminase | Urease | L-Asparaginase | Amidase |
| --- | --- | --- | --- | --- |
|  | (mg NH<sub>4</sub>-N/g dwt/2h) |  |  |  |
| Slash + Burn | 54.54a | 45.21a | 33.25a | -29.75b |
| Slash Only | 2.33c | 16.92ab | 12.25ab | 49.29a |
| Slash + Ridge | 22.46ab | 4.96b | 3.50b | -18.96b |
| Herbicide | 14.29b | 14.58ab | 7.88b | 63.29a |

Mean values in the same column followed by different letters in each year are significantly different from one another based on Tukey's test at $p \leq 0.05$ .

### 3.4 β-glucosidase activity

Figure 1 shows the effect of tillage methods on β-glucosidase activity in the soil. There were significant differences (P ≤ 0.05) in the effects of tillage methods on soil β-glucosidase activity. Slash + ridge (618.00 µg ρ-nitrophenol/g dwt/h) had the highest β-glucosidase activity with significant difference from other tillage methods. This was followed by the herbicide applied plots with a mean value of 519.08 µg ρ-nitrophenol/g dwt/h, which do not differ significantly from slash only plots with a mean value of 493.16 µg ρ-nitrophenol/g dwt/h. Also, slash + burn (298.98 µg ρ-nitrophenol/g dwt/h) recorded the least mean value for β-glucosidase activity which differed significantly (P ≤ 0.05) from other tillage methods.

**Fig 1.**
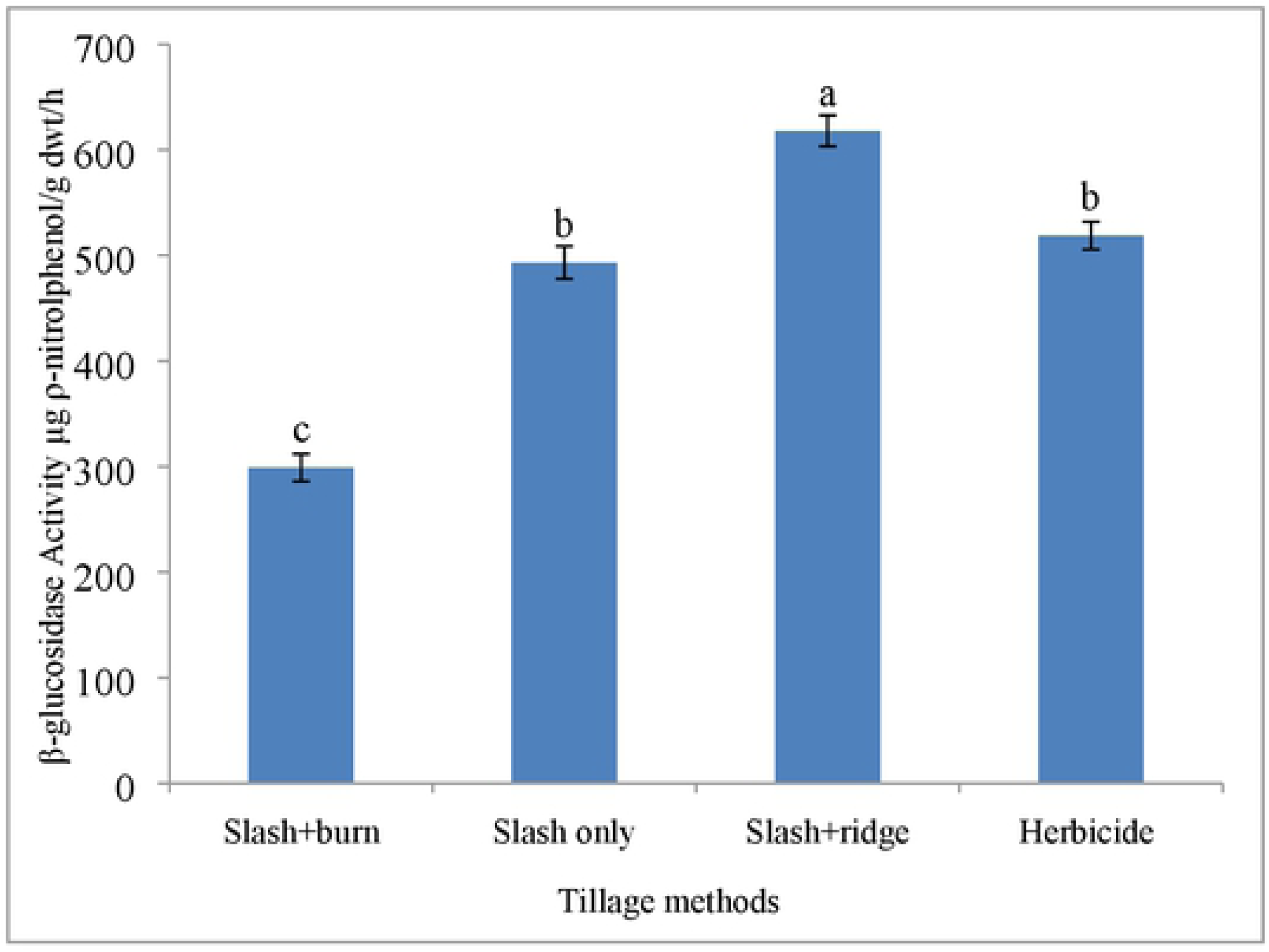
Effects of tillage methods on β-glucosidase activity

### 3.5 Soil dehydrogenase activity

Figure 2 shows the effect of tillage methods on soil dehydrogenase activity. During the study, significant differences existed (P ≤ 0.05) in the effects of tillage methods on soil dehydrogenase activity. Slash + burn (46.11 µg TPF/g dwt/h) recorded the highest mean value for soil dehydrogenase activity with significant difference from other tillage methods. This was followed by the herbicide applied plots with a mean value of 16.11 µg TPF/g dwt/h, which do not differ significantly from slash + ridge plots with a mean value of 21.94 µg TPF/g dwt/h. Also, slash only (11.53 µg TPF/g dwt/h) recorded the least mean value for soil dehydrogenase activity which differed significantly (P ≤ 0.05) from other tillage methods.

**Fig 2.**
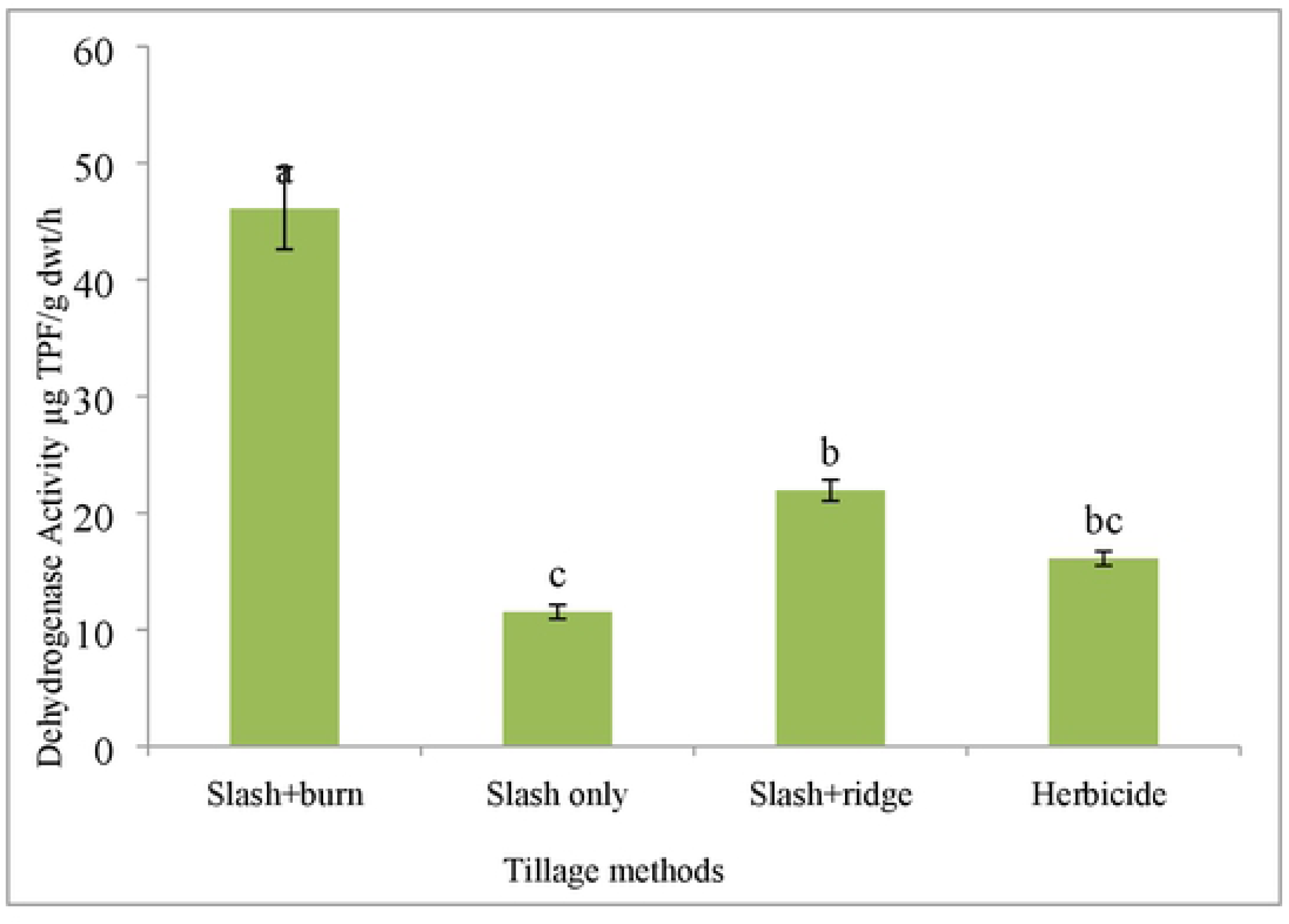
Effects of tillage methods on soil dehydrogenase activity

### 3.6 P-cycling enzyme (phosphomonoesterase) activity

Figure 3 depicts the effects of tillage methods on Phosphomonoesterase activity in the soil. It was observed that tillage methods significantly (P ≤ 0.05) affected soil acid and alkaline phosphate activities. Slash + burn had the highest mean values for acid and alkaline phosphate activities (484.10 and 232.82µg ρ-nitrophenol/g dwt/h), respectively which differed significantly from other tillage methods. This was followed by herbicide application with mean values of 195.61 and 175.10 µg ρ-nitrophenol/g dwt/h, respectively, for acid and alkaline phosphate activities with significant difference from other tillage methods. Slash only recorded least values for both acid and alkaline phosphate activities (59.64 and 40.87 µg ρ-nitrophenol/g dwt/h) respectively.

## 4. Discussions

### 4.1 Soil physico-chemical properties

Conservative tillage practices had, over the years, involved the use of herbicides, slash and burning, ridging, no-till farming, the use of mulch materials, and so on and so forth. These tillage practices, though conservative in nature, had their attendant effects on the soil’s physical and chemical properties. Slash only (SO) practice recorded low growth and yield parameters of maize which might be due to the reduced microbial activity and low water intake of crops, resulting from high bulk density and reduced porosity of the soil. Likewise, Good and Beatty (2011) stated a decline in the cumulative water intake, which they attributed to the soil’s high bulk density and penetrometer resistance under no-till treatment in continuously-cultivated maize with fertilizer. The higher bulk density under no-till method may possibly be as a result of non-integration of organic wastes that remained on the surface of the soil as well as the residual roots of weed which was left in the soil after slashing. Generally, soils with a high bulk density could have impeded root growth and exploit of other plant nutrients; hence, lower yields of maize (Cresswell and Hamilton, 2002). In slash only (SO) tillage plots, decline in yield may also be because there is lack of soil loosening in order to provide favourable conditions for maize growth and yield as compared to better growth and yield in slash + ridge (SR) plots. Findings on maize growth and yield under SO practice further agreed with the results obtained by Ishaq *et al.* (2001). Slash + ridge (SR) gave better yields (about 7% and 12% increase respectively in both years) compared to SO which could be attributed to better soil conditions for seedling establishment and growth, physical modifications of the soil including reduced soil bulk density, increased porosity and water holding capacity, improved nutrient status of the soil (available P, exchangeable K^+^, Ca^2+^ and Mg^2+^) and microbial activity resulting from organic matter incorporation into the soil as a result of ridging. Herbicide (glyphosate) applied plots performed better when compared to SO and SR in all the yield and yield components of maize because the soil available nutrients are utilized maximally owing to little or no weeds-crop plant competition. Also, the killed weeds as well as the applied glyphosate must have served as food source for a wide range of soil animals thereby increasing microbial population, organic matter decomposition and nutrient turn-over rate in the soil. On the other hand, slash + burn (SB) treatment compared favourably amongst all the four tillage treatments in both growth and yield parameters in both years, although, not significantly different from herbicide-treated plots but differed significantly from SR and SO plots. This result is not far-fetched from the fact that nutrient accumulation from years of fallow and release through burning accelerated the growth and yield of maize. In addition, a well-diverse and healthy population of soil microorganisms in SB plots which was stimulated by the prescribed burning might have stabilized the soil ecosystem (Chauhan *et al.,* 2006) from their potential to recycle and restore nutrients for plant growth sustenance. Also, the liming effect of ash produced through burning enhanced the soil pH to a slightly acidic level. At this pH scale, bacterial population and total microbial activity of the soil was improved and most soil nutrients became available.

### 4.2 Soil biochemical properties

The findings from this study have clearly demonstrated that the tillage practices under study including herbicide application can lead to qualitative and quantitative changes in soil enzyme activities. This finding agrees with similar studies from various researchers (Sannino and Gianfreda, 2001; Min et al., 2001; Saeki and Toyota, 2004; Sebiomo et al., 2011). Enzyme activities have been suggested as promising indicators for estimating the extent of contamination in polluted soils because they are observed to be sensitive to chemical pollutants (Aoyama and Nagumo, 1995; Insam et al., 1996; Kuperman and Margret, 1997), and are therefore, referred to as indicators of soil environmental quality (Aon and Colaneri, 2001). Any alternation in soil microbial diversity, their population and activity might function as bioindicator of soil quality and as well reveal its fertility status (Miloševiã et al., 1997; Schloter et al., 2003). Higher soil microbial populations and enzyme activities associated with SB tillage method compared with the other tillage methods may be related to changes in soil water content, pH, organic C and N levels. Soil moisture acts as the prime factor that stimulate microbial populations; soil pH, organic C and N levels might as well considerably influence the activity of microbes as they have been reported as regulators of this parameter (Subhani et al., 2001; Wang et al., 2016). The water contents of the soil, temperature and aeration regimes, and placement of microbial substrates assist in regulating the ecological niches and types of microorganisms which predominate under each of these tillage treatments (Subhani et al., 2001).

The tested activities of β-glucosidase, acid and alkaline phosphatases, as well as the N-cycling enzymes; L-glutaminase, L-asparaginase, amidase and urease have been reported to perform vital roles in the cycles of C, P and N in soils (Anna, 2014). It is notable that the activities of L-glutaminase, L-asparaginase, urease, dehydrogenase, acid and alkaline phosphatases were all enhanced under SB tillage method compared with other tillage methods. No amidase activity was noticeable under SB and SR treatment methods at the end of the growing season, perhaps because amidases have a wide range of substrate specificities (Kobayashi et al. 1993). These observations disagree with those of Hernandez et al. (1997) and Giacomo Certini (2005) who discovered that the activities of urease, alkaline phosphatase, dehydrogenase, arylsulphatase, and N-alpha-benzoyl-L-argininamide hydrolysing protease were reduced by burning alongside soil respiration and microbial biomass C in a Mediterranean pine forest. Boerner et al. (2005) also established that the incidence of prescribed burning reduced the activities of β-glucosidase and acid phosphatase by 5–50% and 15–50%, respectively in a soil under Quercus rubra, which partly agreed with the result of this experiment in the case of β-glucosidase.

Also, the increase in dehydrogenase and phosphatase activities in SB tillage plots agreed with earlier reports that nutrient management practices can lead to an upsurge in soil microbial population and enzymatic activity (Vandana et al., 2012). The increased activity of dehydrogenase could possibly be attributed to a greater availability of soil nutrients usually utilized by enzyme releasing microbes. Unlike enzymes that are associated with specific nutrient mineralization, dehydrogenase activity seemed to rely more on the soil metabolic state and/or on the biological activity of the microbial populace than on any free existing enzyme; hence, situations that stimulate microbial growth are indirectly expected to intensify dehydrogenase activity. Furthermore, the outcomes of earlier microbial studies from this research corroborate with this finding in that it showed higher sensitivity of soil microbial characteristics in SB plots to reflect alterations in management practices and which related favourably with the available soil nutrients (Bandick and Dick, 1999; Bending et al., 2004; Geisseler and Horwath, 2009; Peixoto et al., 2010; André Alves et al., 2013).

A decline in L-glutaminase, dehydrogenase and phosphatase activities were observed in glyphosate treated plots compared to SB tillage plots. This was earlier reported by Min et al., (2001) and Tejada (2009), who separately indicated that herbicides, including glyphosate, stringently inhibited the activity of phosphatases even when applied in dissimilar conditions, with regards to herbicide dosage and soil physical and chemical properties. Beta-glucosidase activity can be ascribed to the labile SOM fraction gradients in the soil and may result in subsequent carbon availability from the remains of decomposed litter and dying microbial biomass (Udawatta et al., 2010). Beta-glucosidase performs crucial function in organic matter decomposition process in the soil derived especially from plant residues. Cellulose degradation by β-glucosidase activity in soils accelerates organic matter decomposition, since in the plant biomass cellulose appears to be of utmost abundance within the polysaccharides family. Besides, the variation in the activity of β-glucosidase enzyme might be a subtle indication to determining organic C richness in the soil (Stott et al., 2010). Additionally, the variations in the soil enzymes activities is consequential of the style of tillage method adopted, thus essentially influencing the cycling of soil nutrients and its fertility maintenance (Paz-Ferreiro et al., 2016).

The findings obtained on the influences of the tillage treatments under study on soil enzyme activities were a representative of the soil fertility status and relative to the growth and yield of maize cultivars, as well as the set of conditions under which the experiment was conducted. However, the reaction of a specific soil enzyme to a given management practice is impulsive for the reason that a particular substrate which inhibits an enzyme in a soil may activate or exert an effect on the same enzyme in a dissimilar soil or on the same soil type in a different locality, as established from the work of Schaffer (1993).

## Conclusion

This study reveals the variations in the physical, chemical and biochemical properties of soil and their implication on soil fertility relative to maize production under different tillage practices. Slash and burn tillage method compared favourably well among all the different tillage treatments in a consistent manner in most of the chemical and biochemical soil quality indicators studied in both cropping trials. Findings from this study have also shown that SB tillage method was beneficial to the soil environment and maize cultivar in that the heat mineralized and released bound nutrients in the soil and residues which contributed to raising the soil pH and enhancing the level of basic cations in the soil, thereby, improving the performance of maize cultivars. This therefore, suggests that controlled fires like slash and burn, prescribed burns, spot burning, stubble burning can serve as a soil management tool as they can be beneficial to the soil microbial ecology and quality, especially to open up a land for maize production following a minimum fallow period of about five years as practiced in this work.

However, further studies to examine whether the increase in enzymes’ activities and microbial indices observed in this research work due to the effects of slash and burn tillage operations would be favourable or destructive for sustainable crop production and soil ecology over a long-term. This perhaps could be done by monitoring specific plant-microbe-soil communications (responses in the rhizosphere and bulk soil).

